# Retention time prediction to facilitate molecular structure identification with tandem mass spectrometry

**DOI:** 10.1101/2022.10.19.512911

**Authors:** Patrik Friedlos, Lilian Gasser, Eliza Harris

**Affiliations:** Student of ETH Zurich, 8092 Zurich, Switzerland; Swiss Data Science Center, ETH Zurich, 8092 Zurich, Switzerland

## Abstract

Comparing measured and predicted chromatographic retention time can improve molecular structure assignment in applications such as coupled liquid chromatography-tandem mass spectrometry. We assess a range of different machine learning methods to predict hydrophobicity, a molecular property that can be used as a proxy for retention time. The performance of the models is evaluated on the benchmark Martel and SAMPL7 datasets. We find that more powerful models perform better when predicting in-sample but not necessarily when generalizing to out-of-sample molecular families. We also find that ensemble methods can outperform individual models. Additionally, a multitask learning model shows promise for improving the generalization ability of graph neural networks for hydrophobicity prediction. Finally, we discuss how the ability of graph neural networks to generalize for molecular property prediction could be improved further.

## 1 Introduction

Identification of molecules in complex mixtures with high performance liquid chromatography followed by tandem mass spectrometry (HPLC-HRMS/MS) has many applications, such as use in clinical laboratories or analysis of pharmaceutical residues [1, 2]. It has been proposed that retention time (RT), measured with HPLC, can be used to improve structural identification with HRMS/MS [3]. For example, one could consider that molecular structures identified by mass spectrometry are false positives if their measured retention time varies greatly from the predicted retention time [4].

When it comes to estimating RT, one can distinguish between look-up, index-based, machine learning and modeling-based methods [5]. Look-up-based methods involve keeping a database of measured RT. Index-based methods estimate the RT of peptides based on the contained amino acids. Modeling-based approaches estimate RT from a physical model of chromatographic separation. Finally, machine learning methods predict RT directly from a machine-readable representation of the chemical structure of the substance. A key advantage of machine learning methods is their intrinsic ability to adapt to new datasets.

A wide range of machine learning algorithms have been considered for RT prediction. Bouwmeester et al. [6] compared the performance of linear models such as Bayesian ridge regression, LASSO regression and linear support vector regression as well as non-linear models such as artificial neural networks, adaptive boosting, gradient boosting, random forests and non-linear support vector regression. Molecular descriptors, such as the count of functional groups, were used as features. They conclude that gradient boosting performs best. However they suggest considering different algorithms as well as blended approaches. More recently, graph neural networks (GNNs) have been found to outperform Bayesian ridge regression, convolutional neural networks and random forests at predicting RT [4].

While mass-to-charge ratios (mz) measured during mass spectrometry are not affected by the setup and conditions of the experiment, RT can vary significantly depending on the experimental setup. Various machine learning methods have been developed to account for changing experimental conditions. Some methods perform well even if the setup-specific training set is small [6]. Other methods use transfer learning to make them applicable in varying setups [4]. Furthermore, one can predict a retention index relative to reference substances or retention order which is more stable across setups than predicting RT itself [7].

An alternative approach is to use machine-learning to predict a physical parameter that is closely related to retention time. The octanol-water partition coefficient (P or logP) measures the hydrophobicity of a substance, which is strongly related to RT [6]. One could hence consider using hydrophobicity as a proxy for predicting RT. Furthermore, hydrophobicity plays an important role in fields such as drug design [8]. Prediction of logP with machine learning methods is a well established field and many different approaches have been developed. Substructure-based methods cut molecules into fragments and estimate their contribution towards logP. Property-based methods predict logP based on other global properties of molecules [9]. Graph neural networks, which were developed more recently, do not require explicit feature selection but implicitly select features from the graph structure of the molecule alone.

Multiple techniques have been successfully applied to logP prediction. With multitask learning, helper tasks are added to the model, such as related properties or predictions of other models, which can lead to increased performance and generalization [10, 11]. Transfer learning approaches leverage easily available data from a related domain to improve model results. For example, one could train a model on a large dataset of predictions from other models and then tilt the model towards a specific molecular space with a smaller dataset [12]. Data augmentation can be used to improve logP prediction, for example by considering all tautomeric forms of a chemical [13]. Improvements have also been made by leveraging predictions of multiple methods and aggregating them into a new model [14]. Furthermore, recent advances in GNNs have been applied to logP prediction [15].

In this report, we assume that estimates of hydrophobicity can be used to estimate RT and improve the performance of tandem mass spectrometry. We considered different machine learning approaches for logP prediction. The models were trained and their performance was evaluated on two independent datasets. Finally, we discuss the results and present some ideas for improvements.

## 2 Methods

We considered a range of different machine learning methods to predict logP, namely multiple linear regression (MLR), random forest (RF), recurrent neural net (RNN) and graph neural network (GNN). Each model was trained five times with the OPERA dataset. We assessed the performance with two benchmarks, the Martel and the SAMPL7 datasets.

### 2.1 Datasets

#### 2.1.1 OPERA

The OPERA dataset contains 13’828 small molecules with measured logP values. The dataset was compiled by Mansouri et al. for the training of the OPERA models. It is based on the PHYSPROP logP dataset, which was further curated. In particular, errors and mismatches in chemical structure formats and identifiers as well as structure validation issues were corrected. The authors then removed low-quality datapoints, outliers and duplicates. The details of this procedure can be found in their paper [16].

The OPERA dataset contains molecules with masses between 26 and 1203 Daltons. The logP values range from -5.08 to 11.29. The distribution of the two metrics is shown in Figure 1. The high quality of the dataset as well as the large range of molecules it covers, at least regarding mass and logP, make it ideal as a training set.

**Figure 1:**
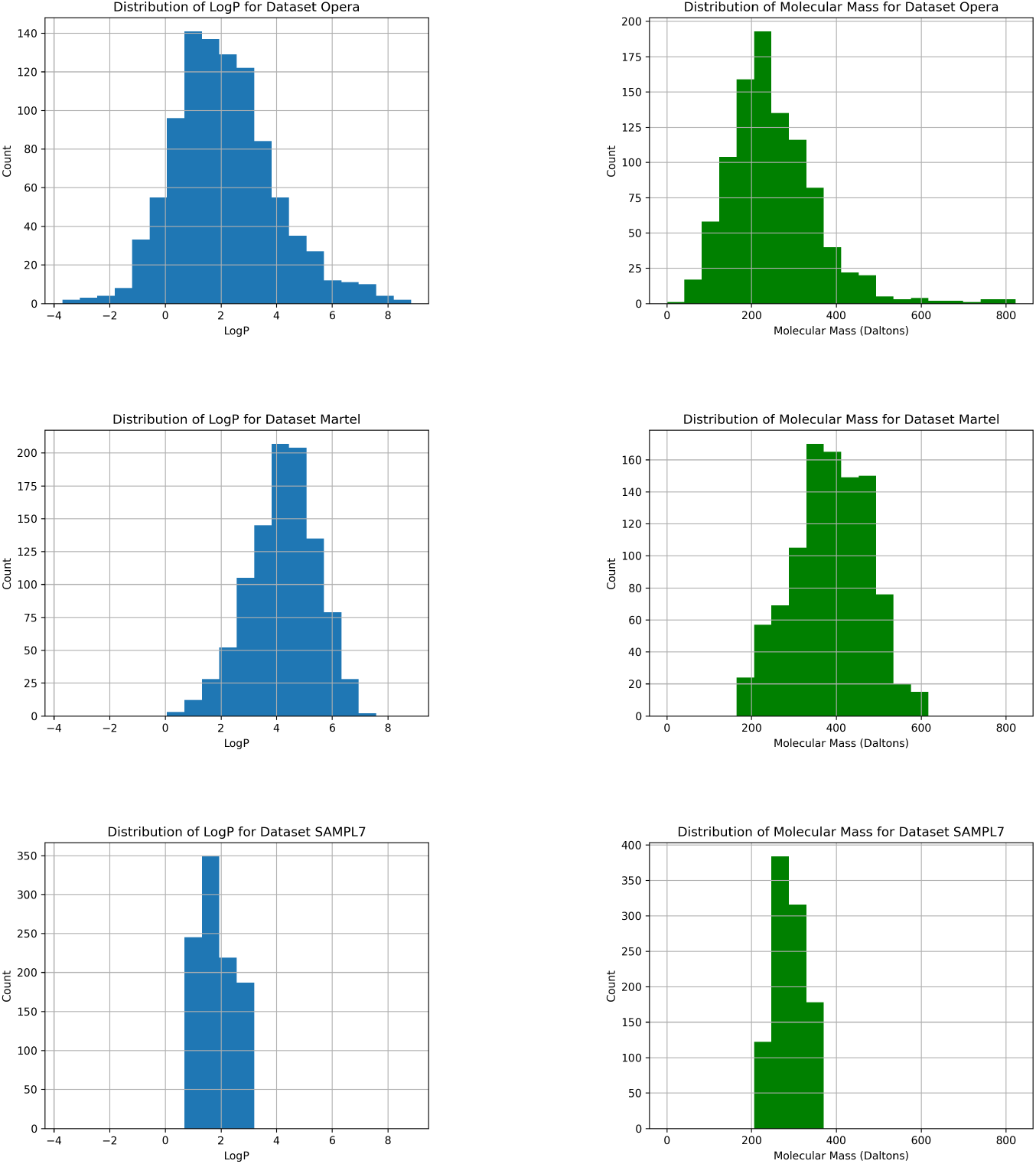
Histograms of logP (left-hand panels) and molecular mass (right-hand panels) for the OPERA, Martel and SAMPL7 datasets. Histograms are shown for the unfiltered datasets.

We removed molecules with a logP above 8, as this report focusses on prediction for small molecules. We further removed molecules that are also contained in the Martel and SAMPL7 datasets to make them viable test sets. A total of 263 molecules were removed, leaving 13565 in the dataset for training.

#### 2.1.2 Martel

The Martel dataset contains logP measurements of chemically diverse molecules for benchmarking studies [17]. The authors constructed a set of diverse molecules, purchased the compounds and experimentally measured the logP values.

The Martel dataset contains 707 molecules with masses between 160 and 599 Daltons. The logP values range from 0.3 to 6.69. The distribution of the two metrics is shown in Figure 1. The Martel dataset is an ideal test set as it is chemically diverse and commonly used as a benchmark, thus results can be compared with previously published studies.

#### 2.1.3 SAMPL7

The SAMPL7 dataset was first used for a blind challenge in physical property prediction [18]. One part of the challenge consists of predicting logP for 22 molecules measured by the authors.

The SAMPL7 dataset contains molecules of masses between 227 and 365 Daltons. The logP values range from 0.76 to 2.96. The distribution of the two metrics is shown in Figure 1. The SAMPL7 is a valuable test set as we can assess the performance of our general-purpose models in a specialized challenge. Some models participating in the challenge were specifically tuned for the molecular space of the challenge [10].

### 2.2 Models

#### 2.2.1 Multiple linear regression

Despite its simplicity, MLR can show good performance in predicting logP. For example, a multiple linear model was among the best performing in the SAMPL7 blind challenge [19]. Similarly, Mannhold et al. proposed a simple linear model based on just the count of carbon atoms and the count of heteroatoms in the molecule. It outperformed many more complex models in their comparison [9].

For the linear model, we need to provide a vectorized summary of molecular structure to use as features. Firstly, we use the count of each atom type contained in the molecule. Secondly, we use all fragment counts available in the RDKit Python package, an open source toolkit for cheminformatics [20]. The regression is then performed using the statsmodels package.

#### 2.2.2 Random forest

We trained a RF non-linear regression model that creates an ensemble of decision trees to predict hydrophobicity. We use the standard implementation of sklearn (RandomForestRegressor from sklearn.ensemble) with default settings. We used the same features as for the MLR model.

#### 2.2.3 Recurrent neural net

While predicting molecular properties with RNNs appears to be less common, they have been used for molecule generation in contexts such as drug discovery [21]. In our RNN implementation, we passed atom representations, as described in the next paragraph, iteratively through the RNN. As there is no natural ordering of atoms in a molecule, we used the default order of RDKit. Finally, we added a fully connected layer predicting logP from the learned representation of the molecule.

The RNN does not require explicit feature generation, however we had to find a suitable representation for each atom within the molecule. We represented each atom as the atom type itself and its neighboring atoms and bond types. We found that there were around 500 different combinations of atoms with their neighboring atoms in our datasets, provided we did not consider hydrogen atoms. We then useed an embedding layer to reduce the dimensionality of the representation.

#### 2.2.4 Graph neural networks

The use of GNNs for logP prediction has been examined by many authors [15]. Atoms are the nodes and bonds the edges of the molecular graph. We used the same atom representation as in the RNN model. After feeding the node representation through two linear layers with a ReLu activation function, six graph convolutional layers as suggested by Kipf and Welling were applied to the graph [22]. The sum was used as the global pooling function. Finally, two more linear layers were used to predict logP. The implementation was done with PyTorch Geometric [23].

In a second GNN model (called GNN + PNA), we wanted to evaluate the benefit of recent advances in neighborhood aggregation for molecular property prediction. In particular, we used Principal Neighbourhood Aggregation (PNA) as suggested by Corso et al. [24]. PNA uses sum, max and var as well as degree-scaled versions thereof in the neighborhood aggregation. We used the Pytorch Geometric implementation of the PNA layer.

In a third GNN model (called GNN + Attention), we wanted to evaluate the findings of Ryu et al. who report benefits of using gated skip-connection and attention for molecular property prediction [25]. To this end, we used a general GNN layer of Pytorch Geometric based on the work of You et al. which features both attention and skip-connections [26].

Finally, we considered a simple GNN with multitask learning (called GNN + Multitask). Multitask learning has been successfully applied to molecular property prediction and logP prediction in particular. Furthermore, it can improve model generalization, which is of major importance should the model be applied to novel molecules. Apart from logP, the model predicts 15 molecular features deemed most relevant by the RF model. This includes counts of atoms but also functional groups, such as the count of benzene rings. The features were computed with the RDKit Python package.

### 2.3 Training

To objectively assess model performances on the Martel and SAMPL7 test sets, we removed these molecules from the OPERA training set. A random 80/20 split of the remaining OPERA dataset was used for training and assessing in-sample performance. When hyperparameters needed to be optimized, an additional validation set was created within the training set.

Every model was trained five times with different random initialization, such that variations in training outcomes can be estimated.

### 2.4 Results

#### 2.4.1 Performance on OPERA

First, we considered the in-sample performance of the models on the OPERA dataset. The root-mean-square error (RMSE) of each model type as well as the standard deviation of the 5 trained models is shown in Table 1. As expected, the more complex models show better performance on both the training and validation set. Most models show signs of overfitting, whereby the performance on the training set is better than on the validation set. The RF shows the strongest overfitting. The MLR does not overfit, which can probably be attributed to model simplicity. Interestingly, the multitask model does not overfit either which could be explained by competition for model bandwidth among the different tasks.

**Table 1:** Comparison of in-sample performance of the models as measured by RMSE for both the training and validation sets using an 80/20 split on the OPERA dataset. The error is the standard deviation due to random initialization.

| Model | RMSE Train | RMSE Test |
| --- | --- | --- |
| MLR | $0.86 \pm 0.00$ | $0.88 \pm 0.00$ |
| RF | $0.25 \pm 0.00$ | $0.61 \pm 0.00$ |
| RNN | $0.27 \pm 0.00$ | $0.45 \pm 0.00$ |
| GNN | $0.27 \pm 0.01$ | $0.45 \pm 0.01$ |
| GNN + PNA | $0.20 \pm 0.01$ | $0.51 \pm 0.01$ |
| GNN + Attention | $0.23 \pm 0.01$ | $0.45 \pm 0.01$ |
| GNN + Multitask | $0.47 \pm 0.01$ | $0.46 \pm 0.00$ |

#### 2.4.2 Performance on Martel

Next, we compared the performance of the models with other methods on the Martel benchmark dataset. Table 2 shows the RMSE of our models and various other models on the Martel dataset. The binned absolute errors of the models can be considered to get a better understanding of the distribution of errors. Table 3 shows the binned absolute errors for our models on the Martel dataset. The values of XlogP3-AA, the best performing non-ensemble model in a study of Plante et al., are added for reference [14]. We can see that the distribution of binned errors is similar between models, however better performing models show fewer large absolute errors.

**Table 2:** RMSE of different models for predictions of hydrophobicity (logP) on the Martel dataset. Results from the models used in this study are highlighted in bold. Unless stated otherwise in a footnote, results are taken from Plante et al. [14]. ^1^COSMO-RS predicts logP with a physical model [13]. ^2^This model uses stochastic gradient descent-optimized multilinear regression with 1438 physicochemical descriptors [27].

| Model | RMSE |
| --- | --- |
| COSMO-RS <sup>1</sup> | 0.93 |
| Lui et al. <sup>2</sup> | 1.03 |
| JLogP-Coeff | 1.04 |
| JLogP—library | 1.05 |
| LogP4Average | 1.12 |
| XlogP3-AA | 1.16 |
| AAM | 1.18 |
| **MLR** | 1.20 ± 0.00 |
| **GNN + Multitask** | 1.20 ± 0.04 |
| **RNN** | 1.21 ± 0.02 |
| Molinspiration | 1.21 |
| SlogP | 1.24 |
| ALogP (Vega) | 1.24 |
| Biobyte CLogP | 1.24 |
| **GNN + Attention** | 1.27 ± 0.05 |
| XLogP2 | 1.28 |
| AlogPS logP | 1.29 |
| **GNN** | 1.30 ± 0.04 |
| **GNN + PNA** | 1.39 ± 0.05 |
| ACD | 1.39 |
| KowWIN | 1.42 |
| **RF** | 1.44 ± 0.01 |
| Meylan (Vega) | 1.66 |
| Mannhold LogP | 1.71 |
| MLogP (Vega) | 1.95 |
| AlogP (CDK) | 3.72 |

**Table 3:** Distribution of absolute errors of the different models for predictions on the Martel dataset.

|  | 0-0.5 | 0.5-1 | 1-1.5 | 1.5-2 | > 2 |
| --- | --- | --- | --- | --- | --- |
| XlogP3-AA | 33.0% | 27.9% | 20.7% | 10.5% | 8.1% |
| MLR | 30.8% | 29.8% | 18.7% | 11.9% | 8.8% |
| RF | 25.1% | 24.4% | 19.7% | 14.9% | 15.8% |
| RNN | 32.8% | 28.1% | 18.3% | 12.0% | 8.8% |
| GNN | 27.9% | 26.4% | 21.4% | 12.7% | 11.6% |
| GNN + PNA | 26.5% | 25.2% | 19.6% | 14.1% | 14.5% |
| GNN + A | 29.3% | 26.8% | 20.5% | 12.6% | 10.8% |
| GNN + M | 32.8% | 29.2% | 18.2% | 10.1% | 9.6% |

We plotted the prediction error as a function of measured logP to assess if the models perform better for some logP ranges. Figure 2 shows the prediction errors of our models against measured logP of the Martel dataset. At least visually, the distribution of prediction errors appears to be homoscedastic. However, the models systematically underestimate logP values for the Martel dataset, with the mismatch increasing at higher logP values. This finding is consistent across different models and can probably be attributed to the differences between the OPERA and the Martel datasets. Whereas the peak of the logP distribution is around a value of 2 for Opera, it is above 4 for the Martel dataset. The models are hence biased towards the logP distribution of the training data.

**Figure 2:**
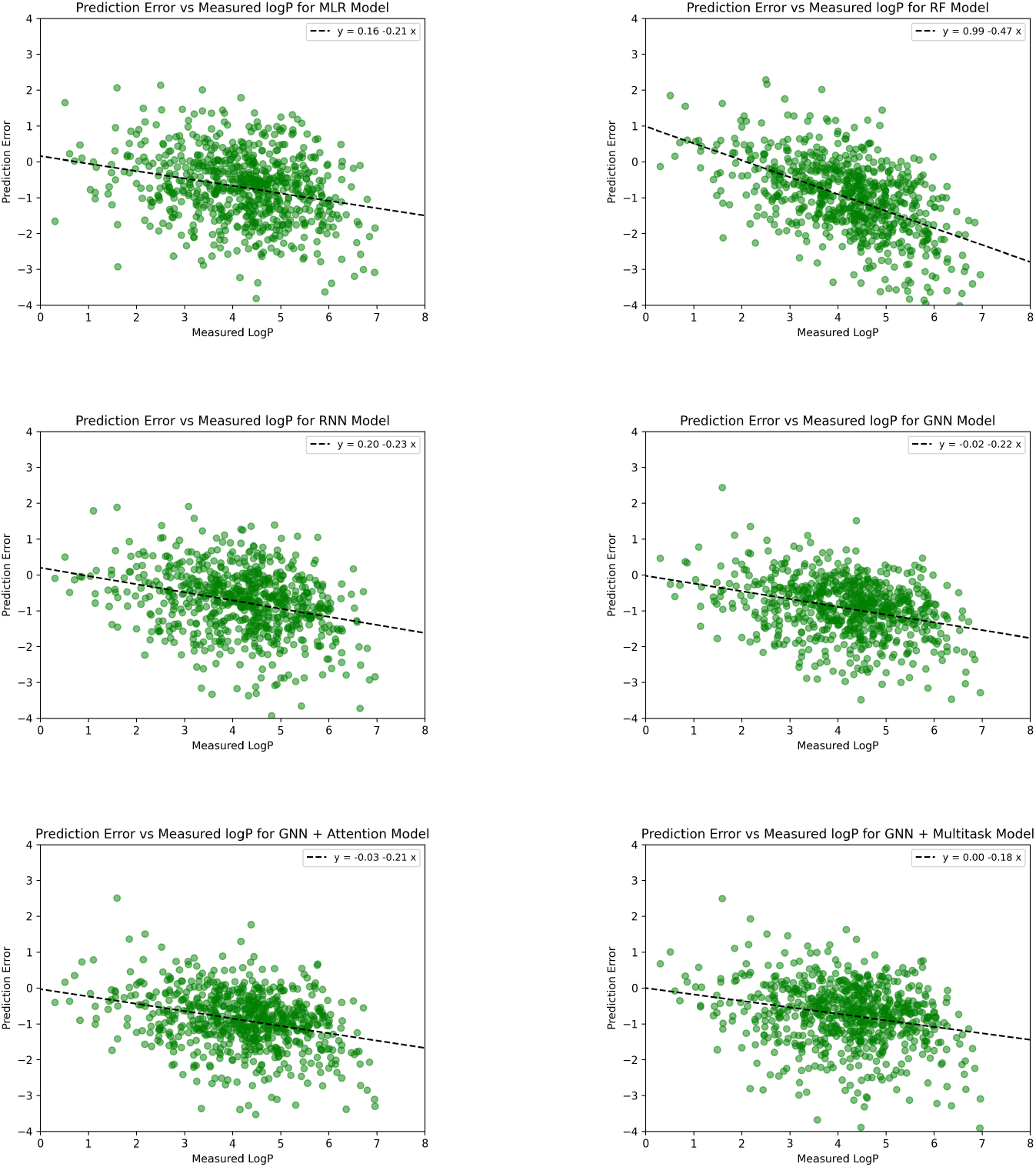
Prediction error compared to measured logP values for each molecule in the Martel dataset. Although the errors appear homoscedastic, the models systematically underestimate logP values as indicated with the linear regression line. All p-values of the regression slopes are less than 0.001.

We considered the Pearson correlation of errors of the different models in order to evaluate to what extent they have learned different information. In case correlations between models errors are low, an ensemble model might be able to outperform the individual models. Table 4 shows the error correlations of the different models on the Martel dataset.

**Table 4:**
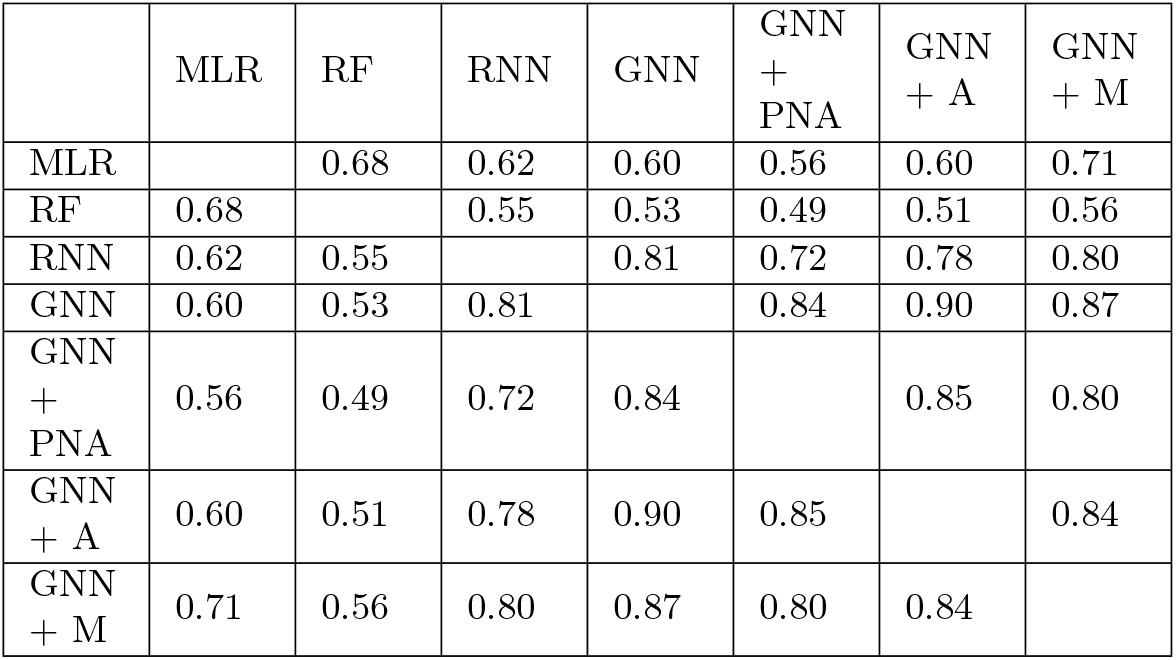
Correlation coefficients between logP prediction errors for the different models on the Martel dataset. The mean error of the five training runs for each model type was used for this comparison.

We can see that model errors on the Martel dataset are correlated between all models. The correlation is particularly high amongst the GNN models, which might be expected, as they belong to the same model family. Interestingly, the errors of the multitask model not only correlate with the other GNNs but also with the MLR and the RF. This might suggest that the multitask model indeed relies on the features learned by the helper tasks to some extent. However, there is no perfect correlation between models and we might expect an ensemble approach to perform even better than any individual model. If we take the mean of all model predictions, the RMSE on Martel turns out to be 1.14, better than any individual model.

#### 2.4.3 Performance on SAMPL7

Finally, we evaluate the performance of the models on the highly specific SAMPL7 dataset. We take all ranked submissions from the SAMPL7 challenge as the benchmark [28]. Table 5 shows the performance of the different approaches on the SAMPL7 dataset as measured by RMSE. We can see that models performing well on Martel don’t necessarily do so on SAMPL7. The GNN + Multitask model however performs well on both test sets.

**Table 5:**
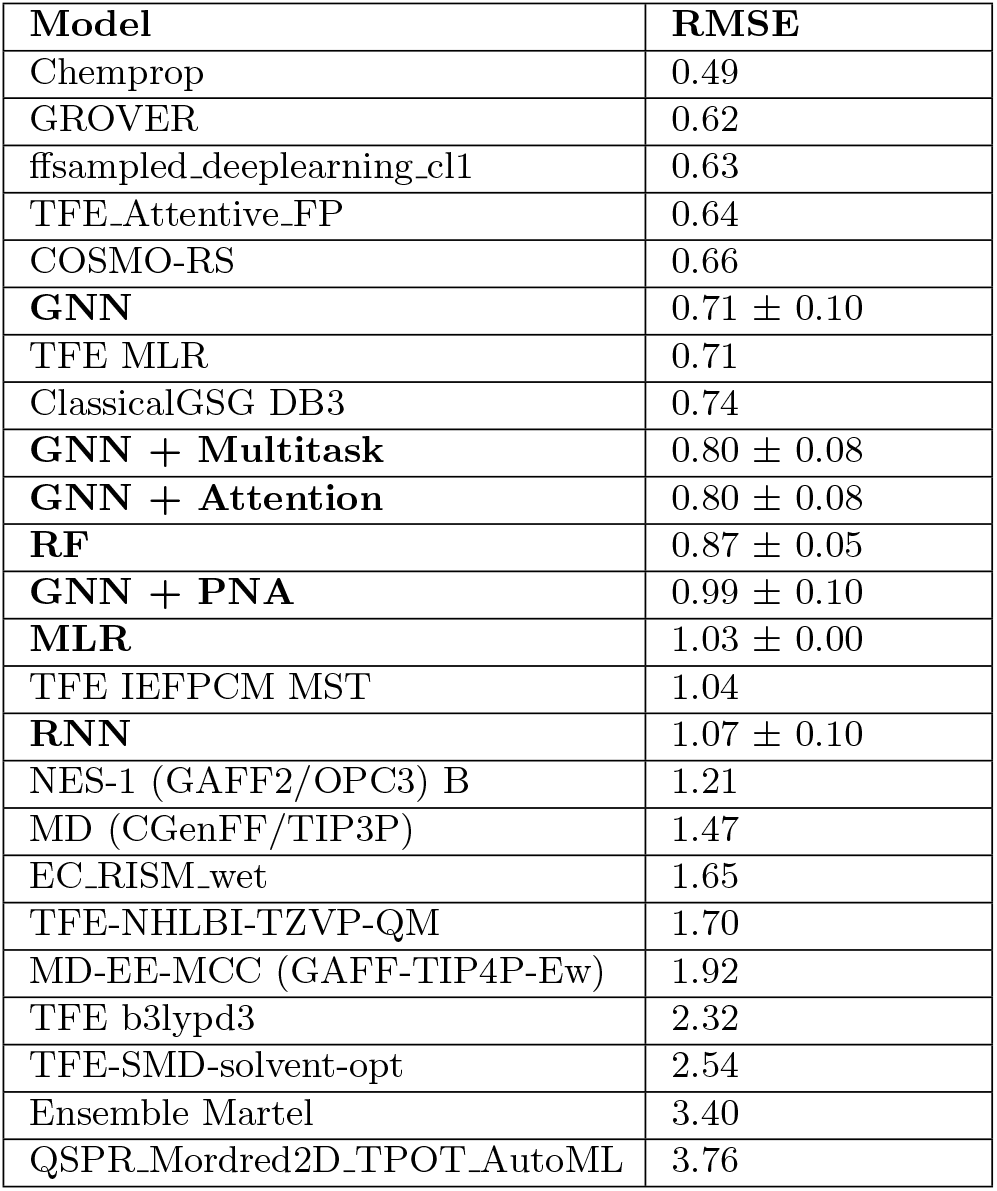
The table shows the performance as measured by RMSE for all ranked submissions to the SAMPL7 challenge compared to our models.

### 2.5 Discussion

#### 2.5.1 Model performance

The results generally show that in-sample performance increases with more powerful modelling methods. As expected, these methods were also more prone to overfitting. The performance of the RNN is remarkable as it performs similarly to the GNNs without complete information on the molecular graph structure. For GNNs we found little benefit in using PNA or attention layers.

For out-of-sample performance on the test datasets, we find that more powerful models offer little benefit a priori. In fact, the MLR performs the best on the Martel dataset despite being the simplest model. Furthermore, there seems to be little correlation between the generalization of the models to Martel and SAMPL7. This could be attributed to the different chemical spaces and logP ranges of the two datasets – the SAMPL7 dataset is more similar to the OPERA training data in terms of logP and molecular weight ranges than the Martel dataset. We also have to consider that SAMPL7 is a small dataset of specific molecules and is not representative for model generalization.

The discrepancy between in-sample and out-of-sample performance can in part be explained by model bias towards the logP value distribution of the in-sample dataset. We have seen that a model trained on the Opera dataset systematically underestimates logP on the Martel dataset. If we train a MLR model with training data from Opera but sampled according to a uniform logP distribution using bins on (-2, 8), RMSE on Martel is reduced from 1.20 to 1.16. If we were to sample according to the logP distribution of Martel, RMSE drops to just 0.96. Of course, the logP distribution of the target dataset can not be known in advance. One could however consider an approach where models are iteratively trained with data sampled according to the logP distribution of the predictions of the previous model.

The multitask model offers the best compromise between in-sample and out-of-sample performance. The loss of performance due to the helper tasks on the in-sample test set is negligible. However, generalization seems to be better, with this model showing good performance on the Martel and SAMPL7 datasets.

All the models considered here performed reasonably well on the SAMPL7 dataset, considering that it contains specific types of molecules. In particular, some models competing in the challenge were specifically tuned for the molecular space.

The variance of model performance is relatively high in particular for the GNNs. This suggests that a significant part of the performance has to be attributed to the initialization. An ensemble of models could be used to improve this issue.

The computational cost of running the models is difficult to estimate, as it depends on the implementation as well as the available hardware. One also has to distinguish between the cost of training the models and using the models. The training cost is typically high for neural nets. Generally an MLR model should be orders of magnitudes faster than a GNN model or a physical models such as COSMO-RS [29]. Predictions can be parallelized however, lowering the runtime of batch predictions if required.

#### 2.5.2 Ideas for model improvement

The implementation of the models was kept relatively straightforward. In particular, we did not perform systematic hyperparameter optimization. For the models requiring explicit feature selection, the MLR and the RF, only atom counts and default fragment counts of RDKit were used. More careful tuning of the models would probably increase performance in some cases.

An interesting paradigm for logP prediction is making it relative to reference molecules with known logP. Such an approach is used by XLOGP3, amongst others [30]. Although this comes with additional challenges, such as having to determine the appropriate reference molecules, it might improve model performance. In particular because such models might not acquire the systematic bias towards logP values found in the training set.

Further improvements could be made by using more datasets for model training and validation. An aggregation of multiple datasets could cover a broader range of molecules and yield increased performance. Different sampling and splitting techniques, such as scaffold splitting [31], could also be considered.

For GNNs specifically, a wide range of different architectures can be considered. This involves choosing from a wide range of convolutional layers, neighborhood aggregations and global pooling layers. Furthermore, there are many options for the initial representation of the atoms. We constructed a representation that includes both the neighboring bonds and atoms. However, there are also setups which handle edge features separately. One could even consider a related graph where every bond in the molecule is a node and the atoms being connected are encoded as node features.

#### 2.5.3 Improving generalization

It seems apparent that a good understanding of the chemical space of both the training set and the target molecules is crucial to achieve good performance on the target set. If they are similar, the use of powerful methods such as GNNs may well be justified. If they are different, simpler models may generalize a lot better. One could imagine that the latter is more often the case in most practical settings.

One solution to achieving good performance on a set of target molecules is the careful construction of a training set. For example, Lenselink et al. created a tailored training set for their participation in the SAMPL7 challenge [10].

Many well-established general logP prediction methods, such as XLOGP3, are essentially linear models [30]. One has to wonder if GNNs could out-perform such methods in broad molecular spaces. The following ideas could be considered to construct a model with broad generalization capabilities:

- Multitask learning is an approach to inductive transfer that improves generalization by using the domain information contained in the training signals of related tasks as an inductive bias [32]. It has already been successfully applied to logP prediction [10]. We have also seen the benefit of this approach, as the GNN + Multitask model showed good performance on validation and test data. As additional tasks, functional groups deemed relevant for logP prediction by other models were used. One could also consider using a RT dataset such as SMRT of METLIN as RT is related to logP [33]. Finally, quantum mechanical properties related to logP could be considered as well. To this end, a dataset such as QMugs could be used [34].
- Transfer learning approaches could be used to enhance performance. In transfer learning, more easily available data from a different but related domain is used for training [35]. For example, a GNN could learn to count functional groups of molecules relevant for logP prediction on a large diverse dataset. The model could then be further trained to predict logP. One would either train a small new part of the model or retrain with elastic weight consolidation to nudge the model to use the more generalizing features learned on the large dataset.
- Combining multitask and transfer learning, a multitask model predicting counts of functional groups and logP could be trained. Next, the model would be used to add predicted logP to a large and diverse dataset for pseudo-rehersal. Finally, the multitask model could be retrained on the large dataset, hoping the helper tasks, on which logP prediction now relies on to a degree, are learned in a more generalizing way.
- It is generally recommended to use a time or scaffold split instead of a random split on the dataset as this provides a more realistic assessment of model performance [10]. One could also tune model parameters with respect to the worst performing of many random splits. Additionally, an adversarial model could be used where an additional model constructs the training set. The resulting model would hence be the best performing on the most difficult split and would arguably generalize better.
- Model generalization to a diverse dataset could be judged a priori to some extent. For example, one could hypothesize that similar distributions of neuron activations between the training and a diverse molecular dataset indicate better generalization. Similarly, a pruning method could be considered in which nodes are removed from the model if its loss reduction on the training set significantly outweighs the loss reduction on the predictions of a large diverse dataset.
- The training set could be augmented to be more representative of a large and diverse dataset. For example, more weight could be added to structurally underrepresented molecules in the training set.

Using any of the above approaches is challenging as they typically come with additional hyperparameters. Not only can the line between too little and too much regularization be narrow, there are also limited independent datasets available for tuning and validation of the approach.

#### 2.5.4 Implications for retention time prediction

It is conceivable that logP predictions can be used as a proxy for RT prediction. However, one should consider that other factors besides logP influence RT, and that the relationship between logP and RT may depend on the chromatography set up. An alternative approach could be a transfer learning approach from existing RT datasets to the specific setup as presented by Yang et al. [4]. The value of using RT in tandem mass spectrometry might be greatly improved by not just predicting RT but also errors. This way, molecules with small expected errors on RT can be identified as false positives more aggressively.

## 3 Conclusion

We set out to evaluate different machine learning algorithms for logP prediction to facilitate molecular structural identification with HPLC-HRMS/MS. In agreement with previous studies, we found that different types of models can show good performance. Both explicit feature construction, such as the use of counts of functional groups, as well implicit feature selection through a GNN are feasible. Further, different models create highly, but not perfectly correlated predictions. This indicates that ensemble methods could further improve predictions.

We conclude that a thorough understanding of the molecular space of both the training as well as the target molecules is essential for logP prediction. Models can show great in-sample performance but lack generalization, and vice versa. More powerful approaches tend to show better in-sample performance. Simpler models, such as MLR, demonstrate less overfitting and can be more suitable for predicting logP out-of-sample.

Modern techniques such as GNNs have been used for logP prediction, often in combination with multitask learning or transfer learning and with a specific set of target molecules. Many of the models geared towards logP prediction in broad molecular spaces, such as XlogP3-AA, have been developed many years ago. We have seen that multitask models might be valuable for creating generalizing GNNs for molecular property prediction. Quite possibly, further improvements could be made with the techniques outlined in the discussion.

## Author Information

This work is the result of a student project of Patrik Friedlos at ETH Zurich supervised by Eliza Harris and Lilian Gasser at the Swiss Data Science Center (SDSC).

## Data and Code Availability Statement

Datasets and code used for this work are openly accessible and can be found at github.com/fpatrik/retention-time.

## Acknowlegement

We thank Nathanäel Perraudin and Luis Salamanca for their detailed feedback on preliminary results.

